# Accelerating in-silico saturation mutagenesis using compressed sensing

**DOI:** 10.1101/2021.11.08.467498

**Authors:** Jacob Schreiber, Surag Nair, Akshay Balsubramani, Anshul Kundaje

**Affiliations:** Department of Genetics, Stanford University

## Abstract

In-silico saturation mutagenesis (ISM) is a popular approach in computational genomics for calculating feature attributions on biological sequences that proceeds by systematically perturbing each position in a sequence and recording the difference in model output. However, this method can be slow because systematically perturbing each position requires performing a number of forward passes proportional to the length of the sequence being examined. In this work, we propose a modification of ISM that leverages the principles of compressed sensing to require only a constant number of forward passes, regardless of sequence length, when applied to models that contain operations with a limited receptive field, such as convolutions. Our method, named Yuzu, can reduce the time that ISM spends in convolution operations by several orders of magnitude and, consequently, Yuzu can speed up ISM on several commonly used architectures in genomics by over an order of magnitude. Notably, we found that Yuzu provides speedups that increase with the complexity of the convolution operation and the length of the sequence being analyzed, suggesting that Yuzu provides large benefits in realistic settings. We have made this tool available at https://github.com/kundajelab/yuzu.

## 1 Introduction

A challenge with using modern machine learning methods in practice is that, frequently, their learned logic for transforming input features into output predictions is opaque and difficult for humans to understand. Accordingly, principled approaches for explaining trained machine learning models have been proposed that, for a given example, assign a numerical value to each feature according to some notion of importance in the resulting prediction. Unsurprisingly, a large number of these feature attribution methods have been proposed, but we have seen three main classes of feature attribution methods: gradient-based [1, 2, 3, 4], path-based [5, 6], and counterfactual- or perturbation-based [7, 8, 9]. These approaches have trade-offs, both in terms of theoretical guarantees and in terms of speed in practice. For example, gradient-based methods generally require one backward pass to explain each output from the model, whereas perturbation-based methods generally require one forward pass to explain each input. However, all three classes have the common goal of assigning to each feature in an example a value that corresponds to the relevance of that feature to the output from the model. When applied in a genomics setting, feature attribution methods are a straightforward approach for identifying the nucleotides, amino acids, and motifs of such, that form the core of biochemical mechanisms or interactions [10, 11, 12, 13].

A simple perturbation-based method, in-silico saturation mutagenesis (ISM), proceeds on biological sequences by constructing mutant sequences that each contain one mutation relative to the reference sequence that attributions are being calculated for. These mutants, along with the reference sequence, are then all run through a model. Unlike the gradient-based methods, this model does not need to be continuous or differentiable. The attribution values are then calculated as the difference in output between the mutant sequences and the original sequences. When the input features are categories, such as nucleotide or amino acid identity, the method has the straightforward interpretation of performing a saturated mutagenesis experiment computationally (hence, the name) [14]. A strength of this method is that the number of forward passes does not depend on the number of output tasks.

In parallel with developments in feature attribution methods, progress has also been made in the field of compressed (or compressive) sensing [15, 16, 17]. This field concerns the replacement of a large number of sparse measurements with a smaller number of dense probes where each probe measures a linear combination of the original, sparse, measurements. For example, rather than performing millions of diagnostic tests, each on one person, one would pool together results such that each pool is made up of multiple individuals and each individual contributes to multiple pools. Through principled pool design, one can achieve perfect recovery of the results that each individual test would have given by only measuring the pools and deconvolving the results given the known pool design, effectively increasing the number of individuals that can be tested with the same resources. Compressed sensing has been used to speed up several algorithms and data collection tools that involve sparse values [18, 19, 20]

An interesting property emerges when ISM is applied to neural networks that contain convolution operations: the difference in the convolution output between the reference sequence and each of the mutants is sparse. This sparsity arises because the convolution operation has a limited receptive field and changes to a single input feature cannot influence the output past that field. Consequently, naive ISM wastes a significant amount of computational time recalculating layer outputs for each mutant sequence that, by definition, must be identical to the layer outputs for the reference sequence. A previous approach, fastISM [9], leveraged this property to justify only recalculating layer outputs that are within the receptive field of the mutation. However, fastISM involves running the same number of convolution operations, albiet restricted to subsets of the sequence the size of the receptive field. Because the number of forward passes remains unchanged, in practice, fastISM requires a large batch size to achieve speedups and, due to implementation details, is usually quite a bit slower than naive ISM when the batch size is small. This is particularly problematic in the interactive exploration setting, where only a small number of sequences are being considered at a time while hypotheses are being developed or models are being debugged.

Here, we describe a method, named Yuzu, that speeds up ISM using two ideas: (1) Yuzu operates on the difference in layer outputs (deltas) between the mutated sequences and the reference sequence and (2) Yuzu uses the principles of compressed sensing to compress these sparse deltas into a compact set of probes that convolutions can efficiently operate on. Because the number of probes depends only on the receptive field of each layer, Yuzu requires a constant number of forward passes regardless of the length of the input sequence, whereas naive ISM requires a number of forward passes linear with the length of the sequence. Together, these properties enable Yuzu to achieve speedups on several model architectures on the scale of an order of magnitude on both a CPU and GPU. Notably, Yuzu significantly outperforms fastISM in the interactive setting for all models, where the number of sequences to be analyzed is small, and either outperforms or is similar in performance to fastISM in the large-batch setting.

## 2 Methods

### 2.1 Compressed sensing

Compressed sensing is an approach for recovering a sparse vector *x* ∈ *R*^*n*^ using a dense vector *y* ∈ *R*^*m*^, where *m << n* and *y* is comprised of linear combinations of *x* created using known “sensing matrix” *A* ∈ *R*^*m,n*^ [21]. Importantly, the columns of *A* should have minimal correlation with each other and so are generally composed of random values from either Gaussian or Bernoulli distributions. A theoretical strength of compressed sensing is one can provably achieve exact reconstruction of *x* from *y* and *A* by solving the optimization problem

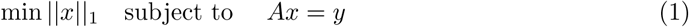

As a note, these results only hold when *x* is *k*-sparse (i.e., has at most *k* non-zero entries) and when *m* ≥ *αk* log *n*, where *α* is some problem-specific constant [21, 15]. These results also extend to when *x* and *y* are both tensors, as will be the case with Yuzu. In the setting where *y* = *Ax* + *ϵ*, where *ϵ* is noise, *x* can still be reconstructed within provable bounds. However, our setting does not involve noise.

### 2.2 In-silico saturated mutagenesis

In-silico saturated mutagenesis is a computational approach used widely in computational biology for calculating the effect that each nucleotide or amino acid has on the predicted output for a given sequence (called the “reference sequence” for the rest of this work). This process involves, first, running the reference sequence through the model and storing the predicted output. Next, sequences are run through the model that each contain a single mutation, i.e., where one position is changed from one character to another. In the context of nucleotide sequences, each mutant contains a single substitution, for a total of 3*L* sequences with *L* being the length of the reference sequence. The attribution score for ISM is then calculated based on aggregating the differences between the outputs when using the original sequence and the outputs when using the mutant sequences. There are several slightly different approaches for using these values to calculate an attribution score for each mutation in the sequence [22, 23, 24], but the simplest is to take the Euclidean distance across all output tasks between the reference and mutant sequence. Finally, these scores are distributed to the features of the original example such that each substitution contains the score that would arise if the feature was mutated, i.e., that the value returned by ISM for a C at a position that is normally an A would be the score calculated from a mutant sequence that contained that A to C mutation.

### 2.3 Compressed in-silico saturated mutagenesis

When ISM is performed on a model that contains convolutions, the differences in layer outputs between the reference sequence and a mutant sequence (the deltas) are limited by the kernel widths and number of the convolution operations in the model, i.e., the receptive field. Specifically, the receptive field for some layer *i* in a convolution-only model is equal to 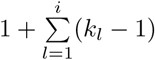 where *l* is a single convolution layer and *k*_*l*_ is the kernel width of that operation. This calculation is more complicated when dilated layers or pooling operations are encountered. Because naive ISM does not account for a limited receptive field, it spends a significant amount of time recomputing values outside the receptive field that must, by definition, be identical to the reference sequence. Calculating ISM attribution scores should only require recalculating the intermediate representations within the receptive field at each layer in the model. An implementation of this idea, fastISM, can achieve significant speed gains (reportedly up to an order of magnitude on common genomics architectures) compared to naive ISM. However, fastISM has a significant initial cost from the bookkeeping associated with a receptive field of increasing size and sequentially applying the convolution operation to a set of 3*L* short sequences.

Our approach, Yuzu, overcomes these costs using a compressed sensing-based approach. A challenge in using compressed sensing theory with neural networks is that most of it only applies to linear models. Although compressed sensing approaches have been developed for non-linear models and do not necessarily require that the output be sparse [25, 26], the guarantees on perfect reconstruction with reasonable computation only apply to linear models [21]. Yuzu side-steps this challenge through sequential application of compressed sensing to each convolution operation individually, because each individual convolution is linear. Specifically, for each layer *ϕ* with inputs from the mutated sequences *x*_*m*_ and inputs from the reference sequence *x*_*r*_, we want to use the layer output from the compact set of probes, *ϕ* (*y*), to reconstruct the difference in layer outputs *ϕ* (*x*_*m*_) − *ϕ* (*x*_*r*_). More formally, we are trying to optimize:

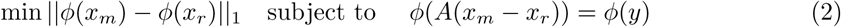

where *y, x*_*m*_, and *x*_*r*_ are tensors, rather than vectors. Notably, even though neither *ϕ* (*x*_*m*_) or *ϕ* (*x*_*r*_) are sparse, *ϕ* (*x*_*m*_) − *ϕ* (*x*_*r*_) is exactly *k*-sparse where *k* is the receptive field of the layer multiplied by the number of potential mutations per position (3 for nucleotides, 19 for amino acids). Because this difference is a linear shift, the optimization problem still holds. Further, because each convolution is a linear operation, we can apply it to both sides such that we build the probes using the layer *inputs* but decode into the layer *outputs*. Conveniently, because we are decoding the difference rather than the outputs separately, we can use the decoded values directly in the next convolution layer to build probes, i.e., the *ϕ* (*x*_*m*_) − *ϕ* (*x*_*r*_) that is decoded in one layer is the *x*_*m*_ −*x*_*r*_ in the next layer, assuming two adjacent convolution operations. As a final note, this optimization is done using a one-stage orthogonal matching pursuit algorithm seeded with the location of the non-zero elements, i.e., linear regression on the non-zero elements.

A key detail is that Yuzu, in contrast to fastISM, operates primarily on the deltas. Yuzu begins by producing a delta tensor that encodes each mutation in a sparse format (Figure 1A). This delta tensor is then passed through the model one layer at a time. Depending on the operation encountered, there are five options that Yuzu can take:

**Figure 1:**
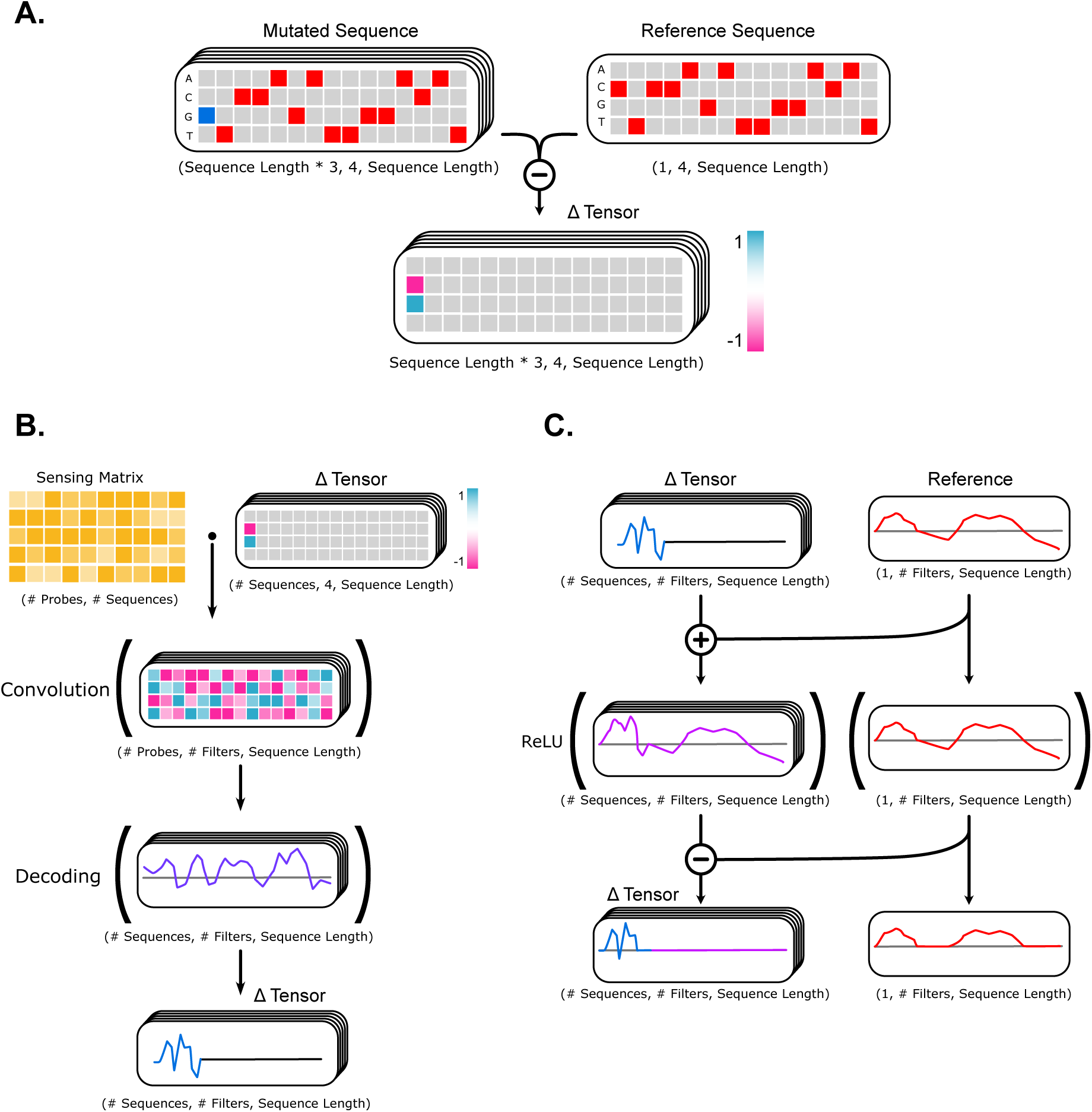
A schematic of the Yuzu process. (A) At the beginning of the process, a tensor of deltas is constructed as the element-wise subtraction of each of the mutated sequences with the reference sequence. (B) When a convolution operation is encountered, probes are constructed as the dot product between the delta tensor and the sensing matrix, the convolution is run on the probes, and the output is decoded back into the signal that would have been observed if the delta tensor originally was run through the convolution. (C) When an element-wise or position-wise operation is encountered, the reference sequence is added to the delta tensor, the operation is performed, and the new reference sequence is subtracted to reconstruct the delta tensor. An important note is that, although the delta tensors in the cartoon appear to have many zeros in them for conceptual simplicity, in practice the delta tensor does not include any columns that are entirely zeroes, and the original delta tensor is constructed directly, rather than as the result of an element-wise subtraction.

1. A convolution layer: the delta tensor is compressed into a set of probes using the sensing matrix associated with the layer (see Section 2.4 for details), the convolution operation is applied to the probes, and the deltas are decoded by solving Equation 2. (Figure 1B).
2. A pooling layer: the delta tensor is added to the reference values and a window is extracted such that the number of extracted positions is equal to number of post-pooled positions that could be affected by the deltas. The pooling operation is then applied and the post-pooled reference values are subtracted out to recover the deltas.
3. An element-wise or position-wise layer: for these layers, Yuzu will add the reference values to the deltas, apply the operation, and then subtract out the post-layer reference values to recover the deltas (Figure 1C).
4. A dense layer preceeded by a convolution layer: Although dense layers do not have a limited receptive field, when the inputs are sparse the number of operations can be limited to only calculate dot products when the vectors are non-zero.
5. A dense layer preceeded by a dense layer: Once the receptive field covers the entire input, Yuzu stops operating on the deltas and just runs the representations through the remainder of the model as normal. Once this happens, Yuzu has the same timings as the naive ISM procedure for the remaining layers but will have a smaller overall time because of the layers that the other steps can be applied to.

At the end of this procedure there are two possible outcomes. The first outcome is that the procedure returns a delta tensor because the receptive field of the model does not extend across the entire input window. Feature attribution values can then be quickly calculated directly from the delta tensor. The second outcome is that the procedure returns the original outputs of the model, rather than the differences from reference. This typically happens after steps 4 or 5 are encountered but can also happen when convolutions with large kernel widths, or max-pooling layers, are used. In this situation, the ISM score is calculated in the same manner that naive ISM is calculated.

Regardless of the outcome, this process replaces the bookkeeping neccessary for fastISM with fast matrix multiplies associated with the construction and decoding of probes. Further, because the constructed probes are the same length as the original layer inputs (but a smaller batch size), the layers can be applied mostly as-is to the probes. In practice, this can be much faster than sequentially applying a layer to a large number of small sequences, even when the total number of operations is the same.

### 2.4 Precomputation

Many aspects of the Yuzu procedure rely on values that are only dependent on the model parameters and sequence length but not the content of the sequence. These aspects include, for each layer in the model, the receptive field, the sensing matrices, the regression coefficients for the optimization step, the locations of the deltas in the sequence, and other statistics. For efficiency, these aspects are precomputed once before running the Yuzu procedure and can subsequently be used for any sequence. Specifically, the sensing matrices and regression coefficients are generated and cached before running the Yuzu procedure, and the receptive field and locations of the deltas within the normal data tensor are empirically calculated by running a single random sequence, and its associated mutant sequences, through the model. Although some of these values could be calculated directly from the model alone, the bookkeepping becomes tedious, particularly when dilatations and pooling layers are involved. Finally, the problem-specific multiplier on the number of probes needed, *α*, is determined by scanning over a pre-defined (or user-specified) range of potential values and choosing the first one that causes Yuzu to match naive ISM’s output on a single random sequence with higher than 1−1^*−*6^ Pearson correlation.

Although the empirical calculation of statistics is conceptually straightforward, three minor modifications must be made to the model in the precomputation step to ensure that the exact receptive field is found and that the internal values do not overflow. First, all pooling layers are converted to sum pooling layers to account for when pooling operations are applied to portions of the sequence that only partially overlap with the receptive field. Second, any activation that has the potential to return zero, such as ReLUs, are ignored to ensure that the correct locations of deltas are found. Third, the weights in the first convolution are changed to be an ascending range across the kernel width and across all filters, and the weights in all subsequent convolutions are changed to be entirely ones. Essentially, the first convolution will identify input mutations and the subsequent layers will then propogate this difference across the layer’s receptive field. Importantly, these changes are not made when calculating the actual ISM scores.

## 3 Results

### 3.1 Compressed sensing speeds up the convolution operation

We began our benchmarking by considering the speedups that our compressed sensing procedure provides to a single convolution operation. These initial evaluations were performed in the idealized setting without including the time needed to communicate values between the CPU and the GPU and discarding the time needed to calculate the feature attribution score: the values represent how fast one would expect Yuzu to speed up a convolution operation within a larger model. We timed Yuzu and the naive ISM approach using convolution operations with an increasing number of filters and kernel sizes applied to sequences of increasing length. In these evaluations, Yuzu and the naive ISM approach were used to calculate attribution scores for 100 randomized reference sequences and the reported time is the minimum total time across 20 runs.

We observed that the speed improvements from Yuzu initially increased with the number of filters until 64 filters, then began to decrease around 128 filters, and then plateaued at 256 filters (Figure 2A). Despite the decrease, we still observed large speedups for all tested filter sizes, with Yuzu being up to 116x faster with 8 filters, in the worst case, to being up to 247x faster with 64 filters, in the best case. These results indicate that Yuzu can drastically decrease the time spent in convolution operations in realistic settings. Interestingly, despite the number of filters, the speedup provided by Yuzu increases with the length of the sequence. This observation is consistent with theory: because the receptive field of the model is constant regardless of filter size or sequence length, the number of probes needed to be constructed remains the same. Hence, the convolution operation only needs to be applied to a constant number of probes that are increasing in length, rather than both longer and more numerous sequences, (i.e., reducing complexity from *O*(*n*^2^) to *O*(*n*)) when calculating ISM scores.

**Figure 2:**
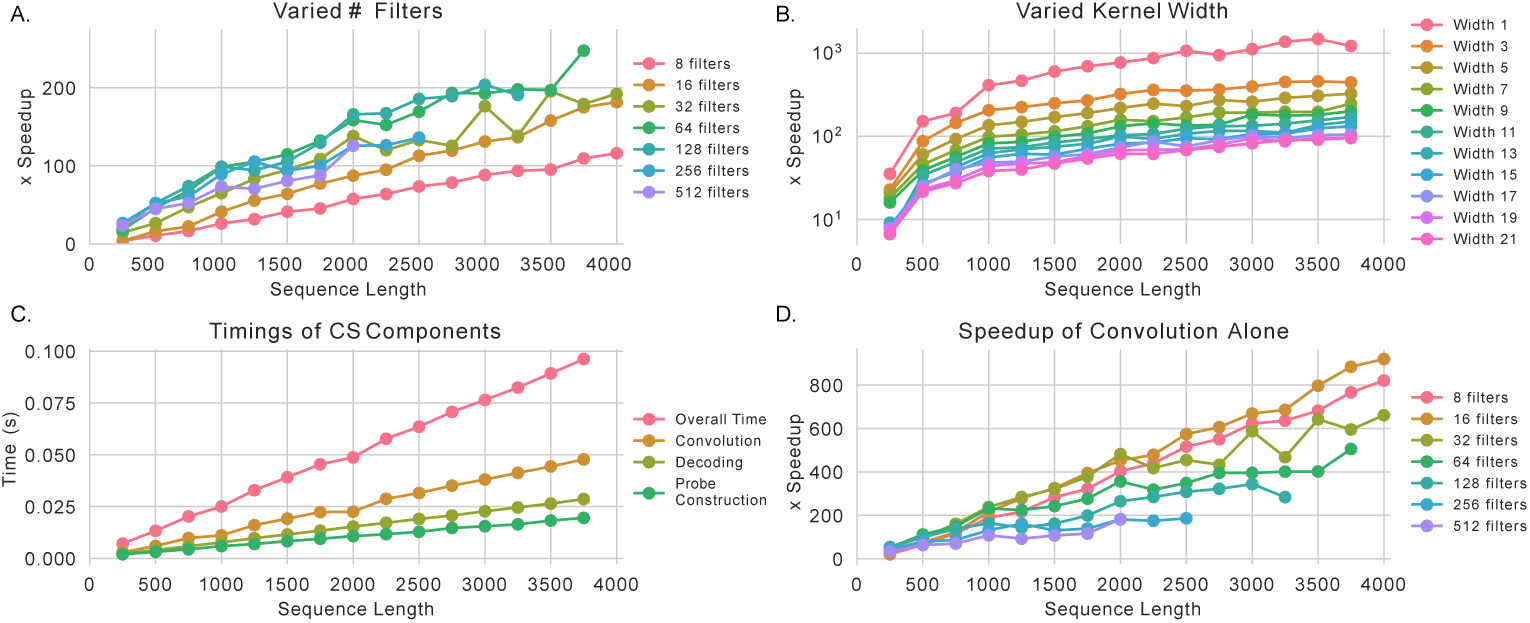
Compressed sensing speeds up convolution operations. (A) The relative speedup of Yuzu when applied to convolutions with an increasing numbers of filters compared to naive ISM applied to convolutions of the same size. (B) The same as (A) except varying the kernel width instead of the number of filters. (C) The total time spent building probes, running the convolution operation on the probes, and decoding the outputs, in the compressed sensing approach. (D) The speedups observed when comparing the time spent running the convolution operation on the probes versus on the entire set of mutated sequences.

Conversely, we observed that increasing the kernel width of the convolution caused decreases in speedups, but that the speedups still did increase as a function of sequence length (Figure 2B). Specifically, a convolution operation with a width of 1 exhibited a 1487x speed increase whereas a convolution operation with a width of 21 only exhibited a 95x speed improvement compared to naive ISM. This finding is also consistent with theory: as the kernel width increases, so too does the receptive field and hence the number of probes that need to be constructed and run through the convolution.

Our next step was to break down the Yuzu timings into the three aspects of the compressed sensing implementation: probe construction, running the convolution, and decoding the output. For a prototypical convolution with 64 filters and a kernel width of 7, we observed that performing the convolution still took a majority of the overall time (Figure 2C). However, the quickness of the probe construction and decoding step speak to the strength of using compressed sensing. Specifically, by spending a small amount of time constructing more informative data, one can greatly decrease the amount of time taken within the convolution while still reconstructing the outputs exactly.

Finally, we compared the time taken by the naive ISM approach, which ran a convolution across all sequences, with the time taken by only the convolution step in Yuzu (Figure 2D). We observed a much cleaner trend with respect to the number of filters, specifically that the more complex the convolution was, the smaller the speedup was. Interesting, this trend was somewhat of a reversal of the original findings. These findings suggest that smaller models find less benefit from the time spent in the probe construction and decoding phases, i.e., that the probe construction and decoding steps take up a greater portion of the overall time and eclipse the benefit from the faster convolution.

### 3.2 Yuzu speeds up ISM on common genomics models

After characterizing the speedups that compressed sensing achieves on individual convolution operations, we next benchmarked Yuzu using several common neural network models used in genomics. Although Yuzu only uses compressed sensing on layers with a limited receptive field, the procedure can still reduce the time spent in most layers because computation is only performed for positions within the receptive field. Specifically, activations, position-wise normalizations such as batch normalization, max-pooling layers, and even the some dense layers can all be sped up in Yuzu (see Section 2.3 for details). Hence, we expected that Yuzu would continue to exhibit good performance even on models with many layers that are not convolutions. In order to achieve a realistic use setting, these evaluations include the time that it takes to produce the mutated sequences and transfer them to the GPU, to perform the inference step (the only part that was timed in the previous set of evaluations), the time it takes to calculate the ISM scores given the outputs from the models, and to transfer the results back to the CPU. Importantly, this evaluation does not include the time necessary to transfer the model to the GPU. Although this time is generally small, we anticipate that both the interactive exploration setting and the large-scale setting would require only a single transfer to the GPU at the beginning before many successive calls of the procedure. The precomputation cost is also not included for the same reason.

Our evaluation involved six pre-defined models: a single convolution, a toy network with three convolutions sandwiching two ReLU activations, DeepSEA, Basset, FactorizedBasset, and a small BPNet model^1^. Rather than changing the complexity of the model or length of the sequence, we evaluated the relative time that Yuzu, naive ISM, and fastISM took when using various batch sizes to calculate feature attributions for 4,096 sequences of length 1kbp and report the minimum time across five runs.

Overall, we observed similar trends across all six models. First, Yuzu exhibited large improvements in speed compared to naive ISM even at batch sizes of 1, whereas fastISM required large batch sizes to outcompete naive ISM. However, although Yuzu and naive ISM both plateaued in speed after certain batch sizes, fastISM exhibited nearly log-linear gains as batch size increased until the batches were too big to fit in GPU memory. For many of these models, particularly the toy models, Yuzu was still faster than fastISM at the largest tested batch size, however, for Basset and BPNet, fastISM exhibited similar speeds at the largest batch sizes. Importantly, the limited receptive field of the OneLayer and ToyNet models meant that Yuzu could calculate ISM scores directly from the final delta tensor without needing to fully reconstruct the output, partially explaining its good performance compared to the other approaches because it did not need to fully reconstruct the output of the model to calculate ISM scores.

Next, we evaluated the effect that changing the number of probes used in the compressed sensing step had on the accuracy of the calculated ISM scores. For each of the six models used before, we varied the number of probes created for each layer through a parameter, *α*, which is a multiplier on the size of the receptive field. At each *α* setting, we calculated the mean-squared-error (MSE) (Figure 3B) and the Pearson correlation (Figure 3C) between the Yuzu feature attributions and the naive ISM feature attributions. As expected, we observed low correlation and high MSE when *α <* 1. However, with only modest values of *α >* 1, Yuzu achieved near-perfect correlation values across all models. Interestingly, DeepSEA appears to achieve reasonable Pearson correlation values even when *α <* 1, but we note that the high MSE suggests that the values themselves are improperly scaled. Overall, we took from this that one needs 1.05 ≤*α*≤1.10. Fortunately, *α* is theoretically independent of the content of the reference sequence and so can be found through a scan of a random sequence in Yuzu’s precompute step.

**Figure 3:**
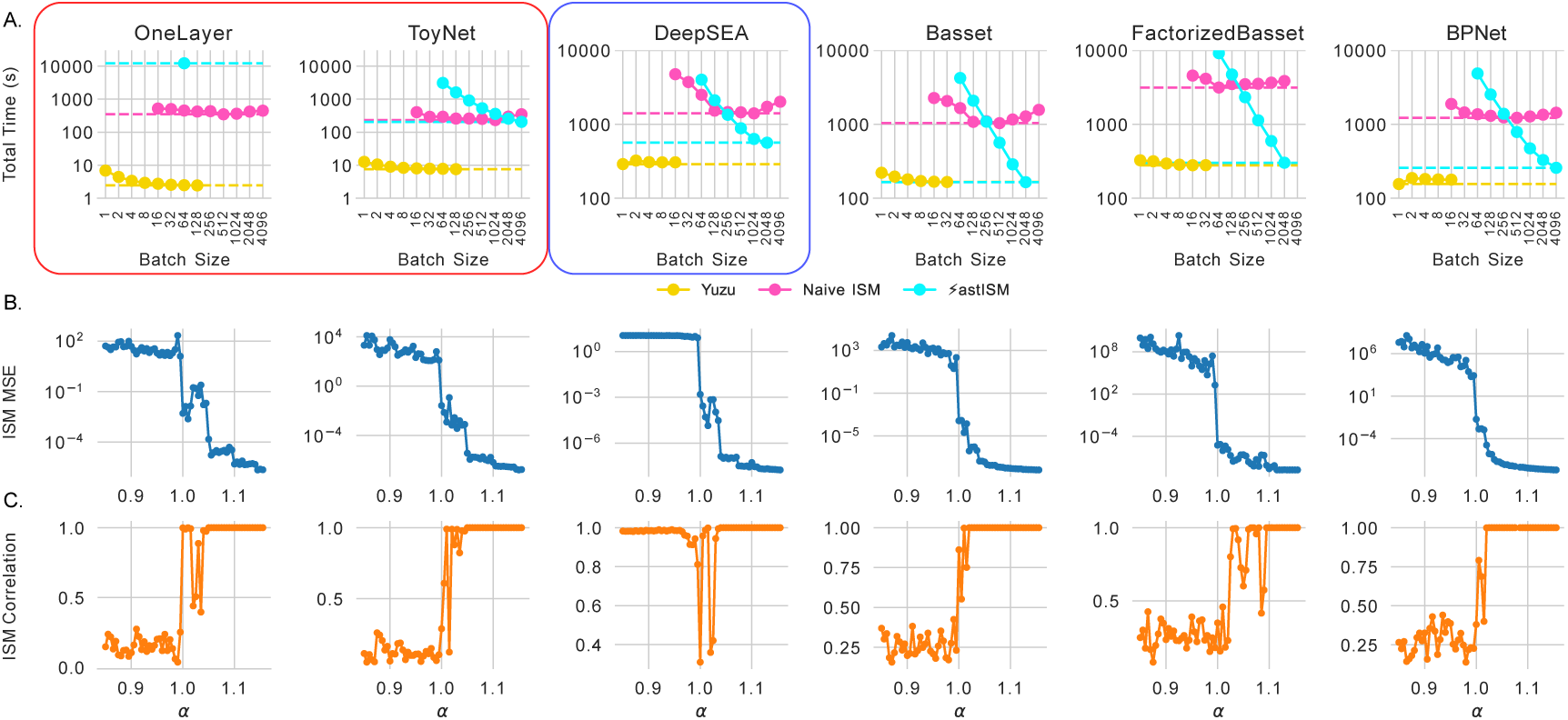
Yuzu speedups on common models. (A) The time spent by Yuzu, naive ISM, and fastISM when calculating attributions for six different models with increasing batch sizes. The dotted line shows the fastest time for each method across all considered batch sizes. The red box indicates models where the receptive field did not cover the entire input window and so Yuzu was able to calculate attributions directly from the delta tensor. The blue box indicates a model where Yuzu was able to use step 4 in it’s procedure without needing to subsequently use step 5. (B) The MSE between the ISM scores produced by Yuzu and those produced by naive ISM as *α* is increased. (C) The same as (B) except the Pearson correlation instead of the MSE.

## 4 Discussion

At a high level, compressed sensing involves replacing a large number of redundant measurements with a smaller number of independent measurements that can be used to reconstruct the redundant measurements. Here, we have demonstrated that compressed sensing can be applied to in-silico saturation mutagenesis by replacing a large number of mutant sequences, whose outputs from convolution layers are largely redundant with the reference sequence, with a smaller number of sequences whose outputs from convolution layers are not redundant with the reference. Empirically, we found that our tool, Yuzu, can speed up individual convolution operations by over three orders of magnitude when compared to naive ISM. When compared with another method with the same goal, fastISM, Yuzu was significantly faster on modest batch sizes but became comparable in speed when massive batch sizes could fit in memory. Together, these results show that Yuzu is most valuable in the interactive exploration setting where one is considering only one or a small number of sequences at a time; however, Yuzu remains competative when applied at scale.

A strength of Yuzu is that it replaces the bookkeeeping necessary for an approach like fastISM with simple matrix multplication to construct probes and decode the outputs. Although a similar number of total operations are performed by fastISM and Yuzu, using compressed sensing allows Yuzu’s implementation to be significantly more compact. As an illustration, the code that is executed when a convolution operation is encountered is only six lines, with three lines spent reshaping tensors. Indeed, one of the largest portions of the code-base is the code executed when a max-pooling layer is encountered, which is very similar to how they are handled in fastISM. Having compact code improves reproducibility and reduces the chances for subtle bugs to exist. Another strength of Yuzu is that improvements in compressed sensing theory can be directly applied to improve its speed even further. For instance, in some contexts one can design a sensing matrix in an informative way instead of using random values [27, 28, 29, 30], potentially requiring far fewer probes to be constructed to achieve perfect reconstruction. Because a large portion of time is spent applying convolution operations to the constructed probes, decreasing the number of probes is a straightforward way to speed up Yuzu further.

Although this work only considers the insertion of a single mutation into each sequence, Yuzu could be extended to the setting where multiple mutations are being inserted at a time. Because the cost of such computation scales with *O*(*n*^*m*+1^) where *m* is the number of mutations inserted per sequence, a similar reduction to *O*(*n*^*m*^) using compressed sensing would be invaluable and could make such analyses more common.

Finally, although the work here primarily involves applications of convolutions to nucleotide sequences, the ideas and the Yuzu tool can be applied more broadly. For example, Yuzu can be applied to amino acid inputs just as easily as nucleotides ones. In that setting, the resulting scores would be analogous to a computational version of deep mutational scanning. Because proteins are short, these types of comprehensive computational analyses can be extremely valuable. Further, the ideas here are not restricted to convolutional operations but can be applied readily to any operation that has a limited receptive field and involves linear operations, such as transformers with sparse attention maps.

## Acknowledgements

We would like to acknowledge Anupama Jha, Yang Lu, and William Stafford Noble for their helpful comments on the paper.

## Footnotes

1 The model architectures follow those proposed in the papers but the weights are initialized randomly, except for BPNet which has only four dilated layers and no residual connections. These changes should not affect the timings of Yuzu, but the random nature of the weights likely requires more probes than weights exhibiting any amount of correlation, as per the restricted isometry property in compressed sensing.

## Notes

### Competing Interest Statement

The authors have declared no competing interest.

